# Adolescent Sugar-Induced Miswiring of Hippocampal Neurons Drives Adult Motivational Dysfunction in rats

**DOI:** 10.64898/2026.10.26.761002

**Authors:** Nuria Masachs, Eve Aurand, Fanny Farrugia, Stephanie Caille, Djoher Nora Abrous, Martine Cador

## Abstract

Adolescence, a critical period for hippocampal maturation, involves the integration of newly generated neurons into circuits underlying adult behavior. While adolescent, but not adult, sugar overconsumption triggers lasting motivational deficits, the underlying mechanisms remain unclear. Here, we demonstrate that adolescent-born dentate gyrus granule neurons causally mediate adult reward-seeking behavior. Strikingly, sucrose overconsumption during adolescence abolishes this function, inducing selective remodeling of synaptic inputs and disrupting their functional integration.

## Main

Adolescence represents a pivotal developmental window^1^ characterized by profound neural plasticity, particularly within the hippocampal dentate gyrus (DG), where newly generated granule neurons (DGNs) are incorporated into existing circuits before neurogenic rates decline in adulthood. These adolescent-born DGNs (Adol-DGNs) are increasingly recognized for their long-term influence on adult behavior^2,3^, including emotional regulation and cognitive flexibility. However, the specific contribution of Adol-DGNs to motivational processes—and their potential vulnerability to environmental stressors such as high-sugar diets—has remained unexplored.

The widespread consumption of sugary diets among adolescents raises critical concerns, as preclinical and clinical evidence links excessive sugar intake to both metabolic dysfunction and adverse mental health outcomes^4^. Notably, sugar overconsumption during adolescence—a period of heightened neuroplasticity—has been shown to induce lasting deficits in adult motivation, emotional processing, and hippocampal neurogenesis, effects that are not observed when sugar overconsumption occurs in adulthood^5^. This suggests that sugar disrupts developmental processes unique to the adolescent brain, potentially altering the maturation and functional integration of Adol-DGNs.

To investigate the involvement of Adol-DGNs in adult motivation, we injected into the DG of adolescent rats, moloney retrovirus (M-RV) encoding inhibitory (Gi) or excitatory (Gs) designer receptors exclusively activated by designer drugs (DREADDs) that allowed us to selectively label and manipulate Adol-DGNs during adulthood^6^.

We first assessed whether inhibiting Adol-DGNs at adulthood would impair motivation for natural rewards. Rats were trained in an operant fixed-ratio (FR) task, where they learned to press a lever to obtain a sweet reward (saccharin). As the fixed ratio increased (from FR1 to FR8), the workload required to obtain the reward escalated, providing a direct measure of motivational drive (time line in **Fig.1a**). Following stabilization of lever performance at FR8 (**Fig.1b**), we administered clozapine-N-oxide (CNO) to inhibit Adol-DGNs and compared lever-pressing behavior to a saline control condition. The results indicated that CNO-induced inhibition of Adol-DGNs significantly reduced the number of lever presses during the first 15 minutes of the session **(Fig.1d-e)**. At the end of the experiment, the recruitment of Adol-DGNs during operant FR8 session was assessed using Egr1 expression as a proxy of neuronal activation, showing that only a small fraction of Adol-DGNs (approx. 2.6%; **Supp.Fig.1**) is activated by the task. These findings demonstrate that Adol-DGNs play a role in sustaining adult reward-seeking behavior.

**Fig. 1:**
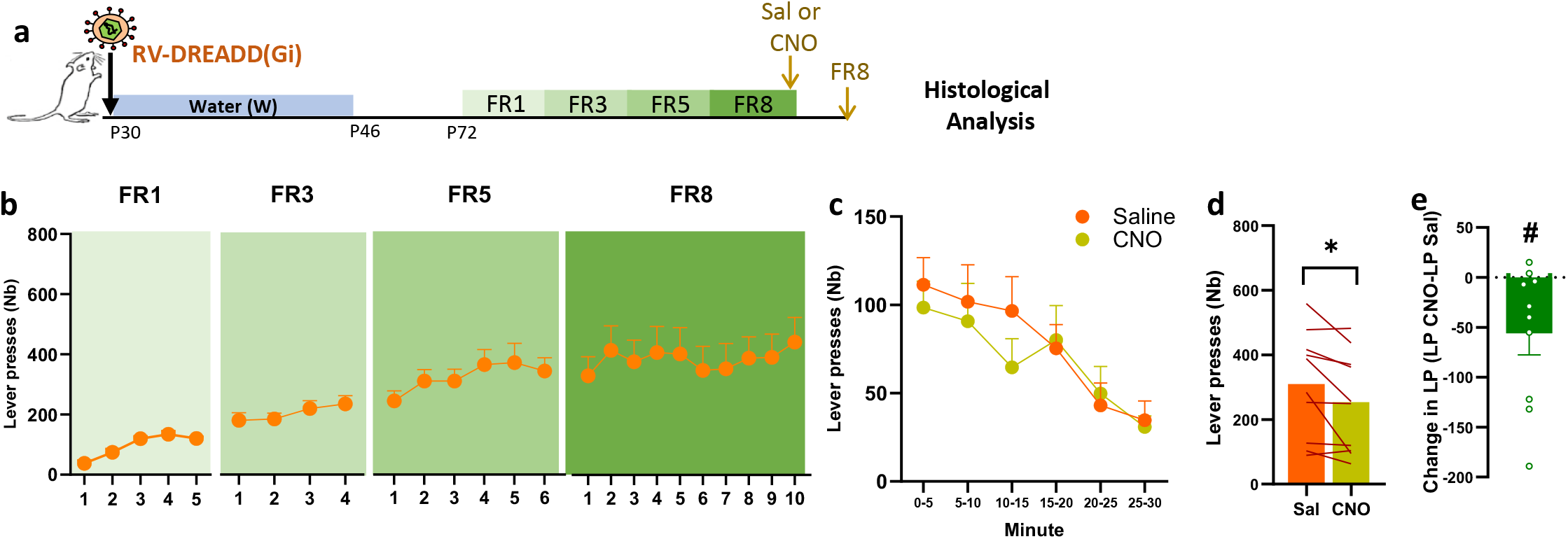
Inhibition of Adol-DGNs controls motivation for saccharin rewards during fixed ratio. **a**.Experimental timeline. P30 rats were injected bilaterally in the dentate gyrus with RV expressing inhibitory (Gi) DREADDs and during 16 days they were isolated and given access to 2 bottles of water (W); once reaching adulthood, animals were trained in operant chambers for saccharin reward under different operant schedules and tested with a saline and a CNO injection in the FR8 schedule. **b.** Evolution of the number of lever presses across the different fixed-ratio (FR) training sessions. **c-e**. After training animals were tested under CNO and saline injections; all animals underwent both CNO and saline sessions. FR8 lever press kinetics by 5 minutes bins (**c**) show no global significant statistical effect of CNO. However, the analysis of the first 15 minutes of the session shows a significant reduction in the number (**d**) and the change (**e**) in lever presses. \**P* < 0.05, # *P<*0.05. Data are mean ± s.e.m. See **Supplementary Table 1** for further details of the statistical analyses.

Next in a mirror experiment, we examined whether **activating** Adol-DGNs could enhance adult motivation and possibly, promote the recovery of motivational deficits in adolescence high-sugar diet exposed animals—known to reduce adult reward-seeking^5^. To test this, as described in **Fig.2a**, a cohort of rats was injected with M-RV excitatory (Gs) DREADDs at P30 and then exposed to either water or 5% sucrose solution from P30 to P46. At adulthood (P72), rats were trained in the same operant FR tasks as above. As expected, sucrose-exposed rats exhibited a significant motivational deficit, pressing the lever less frequently than controls as the workload increased (**Fig.2b**). When Adol-DGNs were chemogenetically activated by CNO, control rats showed a marked increase in lever pressing, confirming that Adol-DGNs play a causal role in motivational processes (**Fig.2c–e**). However, adolescence sucrose-exposed rats failed to exhibit this enhancement, suggesting that sucrose disrupts the functional capacity of Adol-DGNs to support saccharin-seeking behavior. To further challenge the motivational system, we subjected rats to a progressive ratio (PR3) task, where the workload increases exponentially within session, requiring sustained effort to obtain saccharin. Once again, control rats responded robustly to Adol-DGN activation, while sucrose-exposed rats showed no improvement (**Fig.2f–h**). This lack of responsiveness to chemogenetic stimulation in sucrose-exposed animals indicates a lasting impairment in Adol-DGN function, potentially due to structural or synaptic alterations induced by early sugar exposure.

**Fig. 2:**
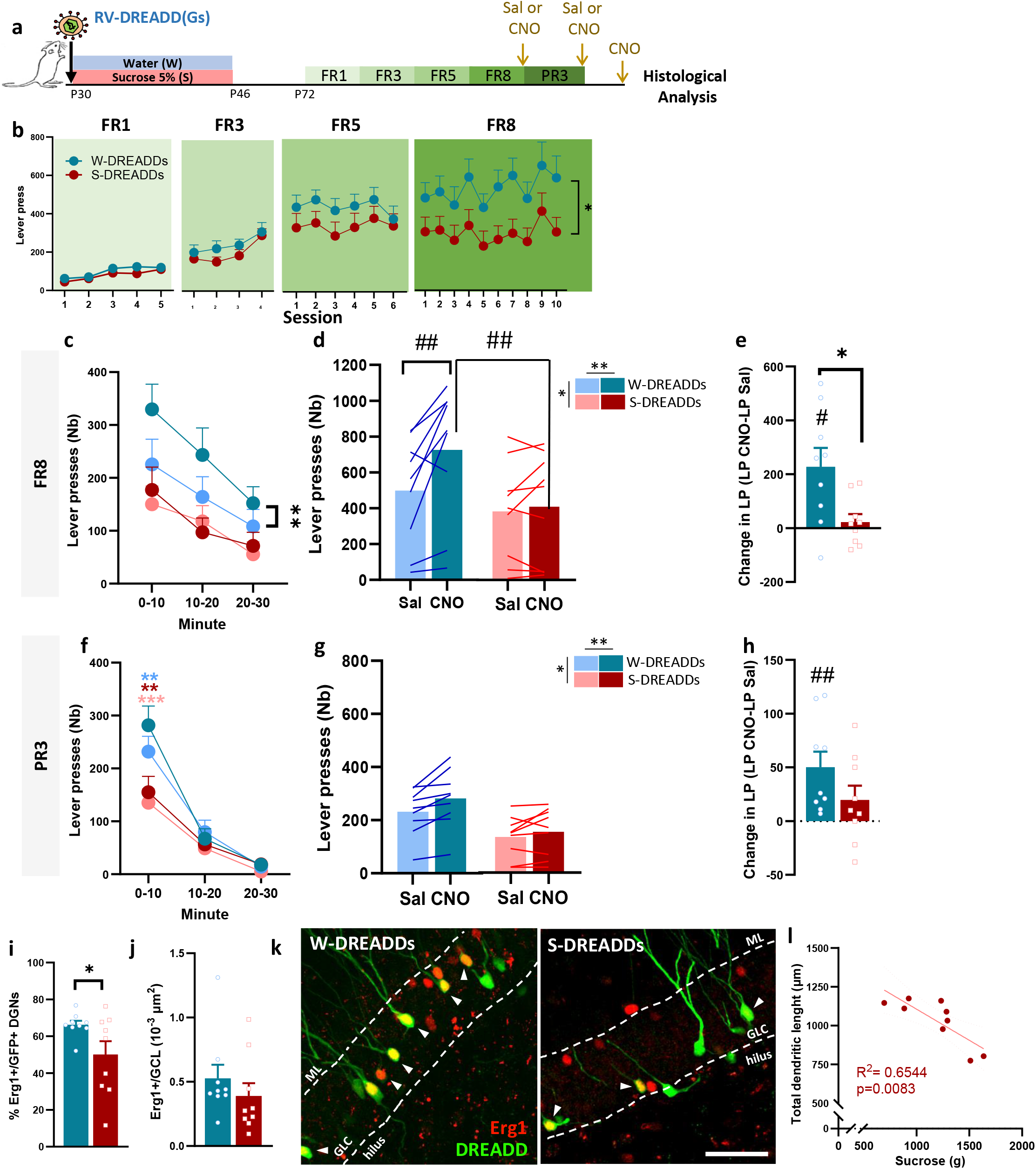
Adol-DGNs regulate motivation for saccharin reward at adulthood and this function is impaired by sucrose overconsumption at adolescence. **a.** Experimental timeline. P30 animals were injected bilaterally in the DG with RV expressing excitatory (Gs) DREADDs and were isolated and given free access to 1 bottle of 5% sucrose and 1 bottle of water (S) or 2 bottles of water (W) during 16 days. Once reaching adulthood, animals were trained in operant chambers for saccharine reward under different operant schedules (FR1, FR3, FR5, FR8) and received an injection of saline or CNO during the FR8 schedule and the PR3 schedule. Following the collection of the behavioral data, brains were processed for histological analyses. **b**. Evolution of the number of lever presses across the different fixed-ratio (FR) training sessions. Sucrose-exposed rats showed blunted responding when FR increased and a stable significant deficit with the highest ratio during FR8 sessions. **c-e**. After training, animals were tested 30 min after receiving a CNO or saline injection (i.p.). All animals underwent both CNO and saline treatment. Kinetics by 10 minutes bins (**c**), total number (**d**) and increase (**e**) in lever presses during FR8 sessions showed that chemogenetic activation of Adol-DGNs markedly increased lever presses in water animals throughout all sessions whereas no effect was detectable in sucrose-exposed rats. In the progressive-ratio (PR3) sessions (**f**-**h**), analysis of the kinetics by 10 minutes bins over 30 minutes (mean duration of the PR3 sessions across all rats) revealed, again, an increase of lever presses induced by CNO only in the water animals during the first 10 minutes of the session. Analysis of these first 10 minutes of the session (**g**) showed a global effect of CNO and sucrose exposure in the number of lever presses, and (h) the increase in lever presses confirms only a significant increase compared to 0 in the Water group, showing that activation of DGNs increases motivation for saccharin only in control animals. **i**. CNO-induced activation of Adol-DGNs, assessed via Egr1 expression, was significantly lower in the sucrose group. This blunted activation was not due to an overall difference in the activity of granule cell layer (**j**). **k**. Representative images of Adol-DGNs expressing DREADDs (GFP in green) and Erg1 (in red) from both groups. White arrows show Erg1+GFP+ DGNs. Scale Bar= 50µm. **l**. Dendritic morphology analysis confirmed a reduced total dendritic length in Adol-DGNs in the sucrose group which correlated negatively with the total sucrose consumed during the last 8 days of adolescence exposure. \**P* < 0.05, \*\**P* < 0.01, \*\*\**P* < 0.001, # *P<*0.05 and ## *P<*0.01. Data are mean ± s.e.m. See **Supplementary Table 2-4** for further details of statistical data.

To confirm that Adol-DGN activation was selectively impaired in sucrose-exposed rats, we assessed neural activity in response to CNO administration using Egr1 expression as a proxy of neuronal activation. While the total number of Egr1+ neurons in the granule cell layer of the DG remained unchanged, we observed a marked reduction in Adol-DGNs expressing Egr1+ in the sucrose group compared to the control one (**Fig.2i–k**). This deficit indicates that sucrose overconsumption specifically disrupts Adol-DGN responsiveness to chemogenetic stimulation, pointing to a fundamental dysfunction in these neurons. Further analysis of dendritic morphology in GFP-labeled Adol-DGNs revealed reduced dendritic length and branching complexity in sucrose-exposed rats (**Supp.Fig.2**). Strikingly, the degree of dendritic atrophy correlated negatively with sucrose intake during the final 8 days of exposure (**Fig.2l, Supp.Fig.3**), suggesting a dose-dependent disruption of neuronal maturation. Together, these findings demonstrate that Adol-DGNs critically participate to adult motivational performance, a function that is abolished by adolescent sugar consumption, likely due to impaired neurodevelopmental processes.

To elucidate the mechanisms underlying the functional deficits observed in sucrose-exposed Adol-DGNs, we analyze their dendritic morphology, synaptic inputs, and mitochondrial integrity. As described in Fig. **3a**, at P30, rats were injected in the DG with M-RV encoding GFP (morphology), PSD95:GFP (glutamatergic synapses), Gephyrin:GFP (GABAergic synapses), or mitodsRed (mitochondria). A subset of animals (batch1) was sacrificed immediately after sucrose exposure (P46) to assess early changes, while another cohort (batch2) was sacrificed at adulthood (P72), when rats were sucrose free and behavioral deficits were evident. At P46, no major differences in dendritic length, branching, or complexity were observed between sucrose- and water-exposed rats (**Fig.3b-e**). However, at P72, sucrose-exposed Adol-DGNs exhibited profound structural deficits, including reduced dendritic length, fewer dendritic ends, and decreased complexity in the Sholl analysis (**Fig.3f–i)**. This set of data indicate an impairment of the late-stage dendritic development of Adol-DGNs. Remarkably, however, the number of GFP Adol-DGNs as well as cell proliferation (Ki67+ cells) was unchanged by sucrose overconsumption (**Supp.Fig.4**).

**Fig. 3:**
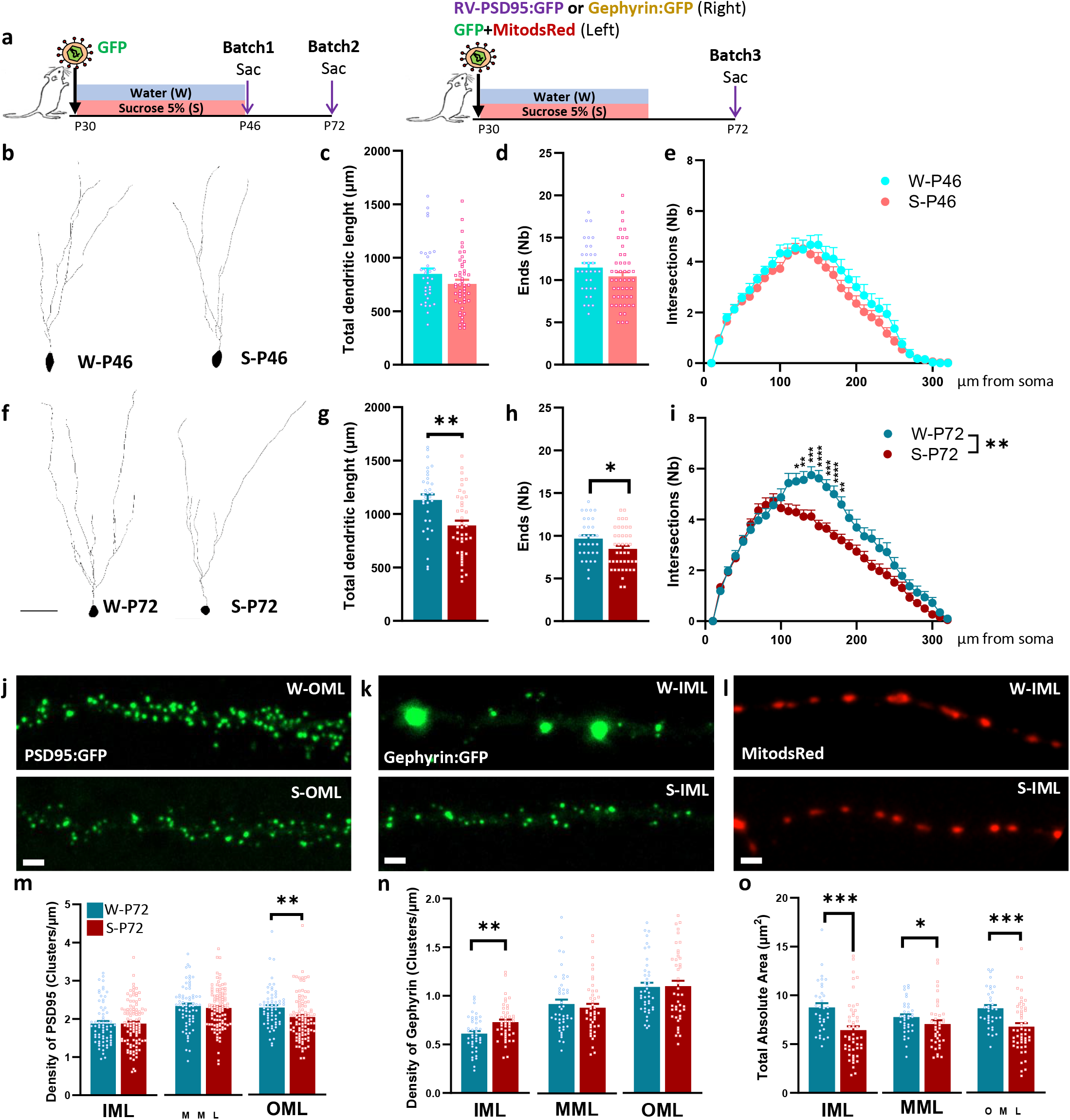
Sucrose overconsumption during adolescence impairs dendritic development, connectivity and mitochondrial network of Adol-DGNs. **a.** Experimental timelines. P30 animals (Batch 1 and 2, left panel) were injected in the dentate gyrus with retrovirus (RV) expressing GFP and were isolated and given free access to 1 bottle of 5% sucrose and 1 bottle of water (S Group) or 2 bottles of water (W group) during 16 days. After this period, batch 1 of rats was sacrificed at P46 and batch 2 at P72 for histological analyses. A third batch of rats (right panel) was injected in the DG with retrovirus expressing GFP + mitodsRed (L: left hemisphere) and PSD95:GFP or Gephyrin (R: right hemisphere) and animals were exposed as previously described and sacrificed (sac) at P72. **b-i**, Representative tracings of Adol-DGNs from water (W) and sucrose (S) group at P46 (b) and P72 (**f**). Scale Bar= 50µm. At P46, sucrose exposure did not alter total dendritic length (**c**), number of ends (**d**) nor the number of intersections in the Sholl analysis (**e**). In contrast, Adol-DGNs at P72 showed clear impairment in dendritic development, evidenced by reduced total dendritic length (**g**), decreased number of ends (**h**) and fewer intersections in the Sholl analysis (**i**) after 16 days sucrose exposure during adolescence. **j-l**. Representative images of PSD95:GFP (**j**), Gephyrin:GFP (**k**) and MitodsRed (**l**) signal at P72 in water (W) and sucrose (S) groups. Scale Bar= 2µm. **m-o**. Analysis of glutamatergic contacts (**m**) at P72 shows a decrease in the cluster density specifically in the OML. In contrast, density of Gephyrin:GFP clusters is affected only in the IML (n) by sucrose overconsumption. Quantification of absolute dendritic mitochondrial area shows a reduction across molecular layer (ML) sublayers – inner (IML), middle (MML) and outer (OML) – in sucrose-exposed animals (**o**). \**P* < 0.05, \*\**P* < 0.01, \*\*\**P* < 0.001 and \*\*\*\**P* < 0.0001. Data are mean ± s.e.m. See **Supplementary Table 9-10** for statistical details.

This neurodevelopmental disruption led us to examine whether sucrose overconsumption alters the synaptic integration of Adol-DGNs by analyzing glutamatergic and GABAergic inputs across the molecular layer (ML) of the DG (Batch 3). In sucrose-exposed rats, glutamatergic inputs (PSD95+) were significantly reduced in the outer ML (OML) (**Fig.3j-m**), a region primarily innervated by cortical inputs. Conversely, GABAergic inputs (Gephyrin+) were increased in the inner ML (IML) (**Fig.3k-n**), and these clusters appeared fragmented and smaller in size (**Supp.Fig.5b**). This shift toward inhibition likely undermines the ability of Adol-DGNs to contribute to hippocampal computations, effectively suppressing their role in reward-related processing.

Given that hippocampal neurogenesis requires adaptation of the mitochondrial compartment to stabilizes dendritic tree as well as to fuel synaptic plasticity^7,8^, together with the evidence that mitochondrial morphology and function is disrupted by high sucrose intake^9^, we further examined mitochondrial organization within Adol-DGNs. We quantified the total area, the average size and number of mitochondrial clusters in P72 Adol-DGNs expressing mitodsRed. In sucrose-exposed Adol-DGNs, we observed a significant reduction in mitochondrial cluster area **(Fig.3l-o)** and size (**Supp.Fig.5c**) across all ML sublayers; despite an unchanged mitochondrial density (**Supp.Fig.5d**). These findings suggest that sucrose modifies the mitochondrial organization, potentially compromising the energetic and plastic capabilities of Adol-DGNs.

In conclusion, our work reveals that Adol-DGNs are essential mediators of adult motivation whose development is profoundly disrupted by adolescent sugar overconsumption. The resulting structural, synaptic, and metabolic alterations effectively “lock” these neurons into a dysfunctional state, rendering them unresponsive to activation and incapable of supporting reward-motivated behavior.

This hypothesis aligns with previous work demonstrating that mitochondrial dysfunction— such as that induced by high sucrose—impairs dendritic plasticity and neuronal excitability in hippocampal DGNS^**8**^. Several studies have shown that glucose and fructose, the components of sucrose, trigger mitochondrial dysfunction at high concentrations^**10**,**11**^, leading to dendritic simplification and reduced synaptic responsiveness. Our findings extend these observations to an in vivo developmental context, revealing how adolescent sugar overconsumption reprograms hippocampal circuits in a manner that persists into adulthood.

The hippocampus, particularly via the ventral subiculum, exerts powerful control over reward-seeking behavior through its projections to the nucleus accumbens^12^ and ventral tegmental area (VTA^13^), both key nodes in the mesolimbic dopamine system and reward related behaviors. By disrupting the maturation and integration of Adol-DGNs, sucrose exposure may dampen hippocampal output to these regions, thereby reducing dopamine neuron responsivity and behavioral vigor^14^.

This study identifies adolescent neurogenesis as one of the sensitive substrate through which environmental factors—such as diet—can durably shape brain function and behavior. The widespread consumption of sugar-rich diets during adolescence, combined with the lasting neural and behavioral consequences observed here, underscores the need to reconsider adolescent nutrition as a critical determinant of mental health across the lifespan.

Future research should explore whether interventions targeting mitochondrial health, synaptic plasticity, or neurogenesis can restore Adol-DGN function and mitigate the long-term effects of early-life sugar exposure. Additionally, investigating the translational relevance of these findings—particularly in the context of human adolescent diet and mental health— could provide valuable insights for preventive and therapeutic strategies. These findings not only advance our understanding of hippocampal circuit maturation but also highlight the urgent need to address adolescent dietary habits as a mean of safeguarding long-term brain health.

## Acknowledgements

The authors gratefully acknowledge Yoan Salafranque (UMR CNRS 5287, University of Bordeaux) for great care of the animals. The authors acknowledge Dr. Fred H. Gage (Salk Institute – La Jolla – CA), Chichung Lie (Friedrich-Alexander-Universität Erlangen-Nürnberg) and Dr. Carlos Lois (California Institute of Technology – Pasadena – CA) and Dr Maria Llorens-Martin for the gift of viral-vectors (GFP, mitodsRed, PSD95-GFP and Gephyrin-GFP). Ultracentrifugations for retrovirus production were done in the Biochemistry and Biophysics Platform of the Bordeaux Neurocampus at the Bordeaux University funded by the LABEX BRAIN (ANR-10-LABX-43).

## Authors’ contribution

NM, DNA and MC designed the experiments and wrote the manuscript. NM performed and analyzed the experiments with the help of EA. FF produced the M-RV. SCG revised the manuscript. DNA and MC conceptualized the project and provided funding.

## Conflict of Interest

Authors declare that they have no competing interests.

### Animals

A total of 78 male Wistar rats (Charles River, Lyon, France) were used. Rats aged 21 days old at the time of delivery were grouped housed (2 rats per cage) in standard cages under an inverted 12/12h light/dark cycle with ad libitum access to food and water. Temperature (22°C) and humidity (60%) were kept constant. Experiments were conducted during the dark phase of the light/dark cycle between 09:00 and 17:00 h.

### Retrovirus production

The M-rv-CAG-GFP has been described previously^1^ and is a gift from Prof. F.H. Gage (Salk Institute, La Jolla, CA). The M-rv-PSD95-GFP has been described previously^2^ and is a gift from Dr C. Lois (Caltech, Pasadena, CA). The M-rv-MitoDsRed has been described previously^3^ and is a gift from Dr D.C. Lie (Friedrich-Alexander Universität Erlangen-Nürnberg, Erlangen, Germany). The M-rv-Gephyrin has been described previously^4^ and is a gift from Dr. M. Llorens-Martin (Universidad Autonoma de Madrid, Spain). The M-rv-Gi-GFP and M-rv-Gs-GFP DREADD were generated by the lab as previously described^5,6^. High retroviruses titres (between 4 × 10^8^ and 2 × 10^10^ Transduction Units/mL) were prepared with a human 293-derived retroviral packaging cell line (293GPG) kindly provided by Dr D.C. Lie. Virus-containing supernatant was harvested three days after transfection with Lipofectamine 2000 (Thermofisher). This supernatant was then cleared from cell debris by centrifugation at 3500 rpm for 15 min and filtered through a 0.45 μm filter (Millipore). Viruses were concentrated by two rounds of centrifugation (19 500 rpm 2 h) and resuspended in PBS

### Stereotaxic injections of Retrovirus

At P30, a retroviral solution was injected bilaterally onto the DG. Rats were anesthetized in a box using a mix of oxygen and 3% isoflurane and maintained asleep with 2% isoflurane during the whole surgery in a stereotaxic apparatus. Animals also received lidocaine (0.1 ml, sub-cutaneous sc, Laocaïne) on the incision site and meloxicam (1ml/kg, sc, Inflacam, Virbac). The retroviral injections were stereotaxically made into the DG with cannulae connected to a Hamilton syringe placed into a micro injector (0.25 μl/min) directly attached to the stereotaxic frame. Cannulae were maintained in position for 3 minutes after the end of the injections to let the suspension infuse. Then the skin was stitched with absorbable sutures and the rat was placed in a recovery chamber (37°C) until it woke up.

For the DREADD experimental cohorts, the M-rv-Gs and M-rv-Gi (2 µl per injection site) were injected bilaterally into two injection sites to tag the dorsal area ([AP]: -2,8mm; [L]:-/+1,5 mm; [P] from skull: - 3,5 mm) and ventral ([AP]: -5 mm; [L]: -/+4,2 mm; [P] from skull: -3.75 mm). For the Adol-DGNs dendritic morphology characterization, M-rv-GFP was injected (1µl). For connectivity the M-rv-PSD95:GFP (1µl) or M-rv-Gephyrin:GFP (1µl) were used. For mitochondrial analysis, a retroviral mix of half M-rv-GFP and half M-rv-MitoDsRed was prepared before injection^7^ (2 µl). All retrovirus were injected in the dorsal dentate gyrus ([AP]: -2,8mm; [L]:-/+1,5 mm; [P] from skull: -3,5 mm), respectively. At the end of the experiment, only rats with labelled neurons in both hemispheres were kept in the analysis.

### Sucrose overconsumption

After surgery, adolescent rats were housed in individual cages and given continuous free-choice for the sucrose group (S group) between a bottle of water and a bottle of 5% sucrose (VWR) from PND30– PND46 as previously described^8^. The control group had access to two bottles of water (W group). Fluid consumption, food intake and body weight were measured every other day. The position of the bottles was alternated every day to control for position preference. Following 16 days of sucrose consumption, all animals were given access to only water in collective cages until the end of the experiments.

### Operant behavior

Once rats reached adulthood, motivation for saccharin was studied in 30 min daily sessions (5 sessions per week) in operant chambers (Imetronic, Pessac, France) equipped with two levers, as previously reported^8^. Adult rats (P72) previously injected with M-rv-Gs or M-rv-Gi were trained to press one of two levers (the active lever) under a fixed-ratio 1 schedule (FR1) of reinforcement (i.e., every active lever press was reinforced by fluid delivery in a dipper situated between the two levers) to obtain 0.1 ml of saccharin solution (0.13%, w/v; Sigma-Aldrich) delivered in 3 s. The delivery of saccharin was paired with a 4 s cue light located above the active lever during which additional lever presses did not result in fluid delivery (time out period). Responses on the other lever (the inactive lever) were recorded as a measure of non-specific activity. The position of the active and inactive levers was counterbalanced between rats. The rats were first tested on FR1 (5 sessions) and then shifted to a FR3 (3 lever presses resulted in fluid delivery, 4 sessions), FR5 (5 lever presses resulted in fluid delivery, 6-7 session) and up to FR8 (8 lever presses to obtain a reward, 10 sessions). The criterion for acquisition of saccharin was a minimum of 50 reinforcements per session over 2 consecutive sessions in the FR1 schedule. The number of lever presses and doses were recorded in each session. Following FR8 training, rats were tested for motivation within a session using a PR3 schedule. In this test, the number of required lever presses increased by 3 for each successive reward, starting at 1 (1, 4, 7, 10, etc.). The number of lever presses and rewards received were collected in each session and considered as a measure of motivation.

### Chemogenetic manipulation of Adol-DGNs

The Dreadd ligand CNO (Clozapin-n-Oxyde, Enzo Life Sciences, #BML-NS105) was dissolved in a saline solution and delivered with one i.p. injection of 1 mg/kg in rats 1h before the test as previously described^6^. Chemogenetic manipulation of Adol-DGNs was performed during FR8 and PR3. These tests were performed across two days, during which rats were randomly assigned to receive either a saline or a CNO injection on the first test day, and the reverse the second day.

### Immunohistochemistry

Animals injected with M-rv-CAG-GFP, M-rv-MitoDsRed and M-rv-PSD95-GFP were perfused transcardially with a phosphate-buffered solution of 4% paraformaldehyde at the end of the experiment. After 24h fixation, brains were transferred in a PBS-sucrose 30% solution for 24h and then cut using a vibratome. Animals injected with M-rv-Gs rats were perfused transcardially 120 minutes after CNO injection (Gs). M-rv-Gi rats were perfused 90 minutes after behaviour. After 1 week of fixation, brains were cut using a vibratome.

The activation of Gs-GFP+ and Gi-GFP+ Adol-DGNs was studied using immunohistofluorescence procedure on alternate one-in-eight sections. The retrovirus-expressing GFP was visualized with an anti-GFP primary antibody (Chicken, 1:1000; Abcam, Cat#ab13970) and neuronal activation with an anti-Erg1 antibody (Rabbit, 1:500; Cell Signalling, Cat#4153S). Bound antibodies were visualized with Alexa488 goat anti-chicken (1:1000; Thermo Fisher, Cat#A-11039) and Alexa-568 goat anti-rabbit (Life technologies, 1:1000, Cat#11036) secondary antibodies. Primary antibodies were incubated simultaneously at 4 °C for 48-72h, and secondary antibodies were incubated simultaneously at room temperature for 2h. Fluorescence was studied using a Zeiss LSM900 confocal system with X 20 lens and a digital zoom of 1.5. The percentage of GFP cells expressing Erg1 (all along the temporal–septal axis) was calculated as follows: (Nb of GFP+-Erg1+ cells)/(Nb of GFP+-IEG™ cells + Nb of GFP+-Erg1+ cells) × 100. In all experiments, a minimum of 100 Adol-DGNs were analyzed per rat (**Supplementary table 4 for details**). Gs-GFP+ neurons were also used for the morphological analysis. Images were obtained with a Zeiss LSM900 confocal system with X20 lens (tile module option). Confocal stacks were analysed using ImageJ plug-in SNT to obtain dendritic parameters as well as Sholl analysis. Data shown are from 9 rats per group with a minimum of 5 neurons per animal.

For Ki67 and GFP counting, expressing cells from free-floating 50-μm-thick sections were processed according to a standard immunohistochemical procedure on alternate one-in-eight sections. Primary antibodies used were anti-Ki67 (Mouse, 1:100; Leica Biosystems; NCL-L-Ki67) and anti-GFP primary antibody (Rabbit, 1:1000; Millipore, Cat#AB3080P), followed by incubation with biotinylated goat anti-mouse antibody (1:200; Vector Labs, Cat#BA-2000) or goat anti-rabbit antibody (1:200; Jackson ImmunoResearch, Cat#111-035-144). Bound antibodies were visualized using the biotin-streptavidin technique (Ultra-Sensitive ABC kit, Thermofisher, Cat#32050) and 3,3’-diaminobenzidine as a chromogen (Sigma-Aldrich). Ki67+ and GFP+ cells in all the experiments were counted under a x20 microscope objective throughout the entire septotemporal axis of the granule layer of the dentate gyrus. The total number of cells was estimated using the optical fractionator method, and the resulting numbers were tallied and multiplied by the inverse of the sections sampling fraction (1/ssf8) **(GFP results in Supplementary Table 7**). The morphometric analysis of virus-labelled neurons was performed with a ×100 objective. Measurements of dendritic parameters as well as Sholl analysis were performed with the Neurolucida (software Microbrightfield, Colechester, VT, USA). The analysis of the dendritic arborisation was performed as previously described^9^. Data shown is from 4-6 rats per group with a minimum of 8 neurons per animal.

PSD95-GFP and Gephyrin:GFP were visualized with an anti-GFP primary antiboby (Rabbit, 1:1000; Millipore, Cat#AB3080P). GFP was visualized with an anti-GFP primary antibody (Chicken, 1:1000; Abcam, Cat#ab13970) and anti-DsRed primary antibody (Rabbit, 1:1000; Clontech Takara, Cat#632496). Bound antibodies were visualized with FITC goat anti-Rabbit (1:1000; Jackson Immuno Research, #111-095-003) Alexa-488 goat anti-chicken (1:1000; Thermofisher, Cat#A11039) and Alexa-568 goat anti-rabbit (1:1000; Life technologies, Cat#11036) secondary antibody. Primary antibodies were incubated simultaneously at 4°C for 24-48h, and secondary antibodies were incubated simultaneously at room temperature (RT) for 2h. Analysis of PSD95, Gephyrin and MitodsRed were performed as previously described^4,7^. Fluorescence was studied using a Zeiss LSM900 confocal system with a X 63 oil lens (Plan apochromat Zeiss, numerical aperture 1.4) and a digital zoom of 1 (MitodsRed) and 1.5 (PSD95 and Gephyrin). For both approaches, MosaicJ plugin from ImageJ was used to reconstruct the neuron as a whole. Mosaic reconstruction allowed to determine the length of the molecular layer and to delimitate the inner (IML), middle (MML) and outer (OML) sub-layers of the molecular layer. For each layer, segments of 30 µm were analysed. The number of clusters of PSD95-GFP and the cluster’s density, average size of the cluster and total absolute area and number of mitoDsRed and Gephyrin from the segments were determined semi-automatically using an intensity threshold in Image J. The density of clusters was obtained by dividing the number of clusters by the length of the corresponding dendritic segment.

### Statistical Analysis

The data (mean ± s.e.m.) were analyzed using one sample t-test to compare to 0, t-test (unpaired and paired), 2- and 3-way repeated measures ANOVAs, which were followed by Bonferroni post-hoc tests whenever interactions were present.

