## supplementary data for "Adolescent Sugar-Induced Miswiring of Hippocampal Neurons Drives Adult Motivational Dysfunction in rats"

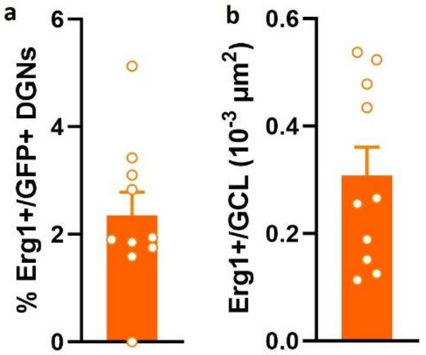


**Supplementary Fig. 1: Adol-DGNs are recruited during FR8 session.**

a. Adol-DGNs recruitment, assessed via Erg1 expression, after a FR8 session. b. Total expression of Erg1+ cells in the granule cell layer (GCL) of DG.

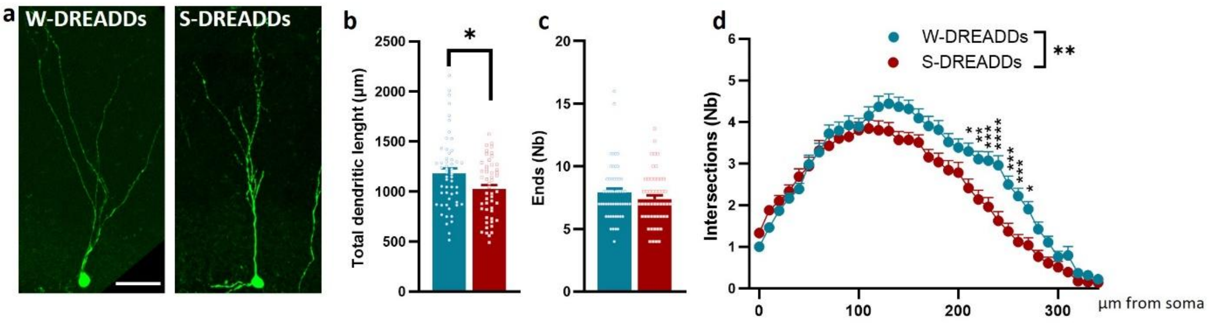


**Supplementary Fig. 2: Sucrose overconsumption during adolescence alters dendritic morphology in Gs-DREADD expressing Adol-DGNs.**

a. Representative images of Adol-DGNs from water (W) and sucrose (S) group. Scale Bar= 50µm. Analysis of Gs-DREADDs Adol-DGNs showed a reduced total dendritic length (b) and fewer intersections in the Sholl analysis (d), however, no significant decrease in the number of ends (c) (n= 54 and 51 neurons from 9 animals per group for W-P72 and S-P72 groups, respectively). *$P < 0.05$, **$P < 0.01$, ***$P < 0.001$, ****$P < 0.0001$. Data are mean ± s.e.m. See Supplementary Table 5 for further details of statistical analyses.

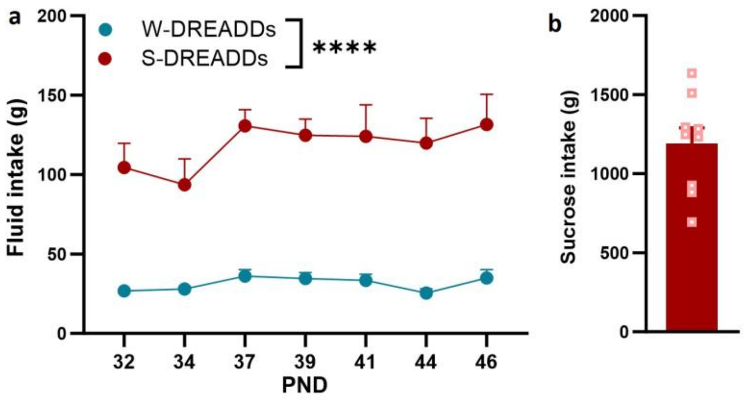


**Supplementary Fig. 3: Ad libitum access to 5% sucrose during adolescence strongly increased fluid intake**

a. Fluid intake from PND30 to PND46 of Water and Sucrose rats. b. Total sucrose intake during the last 8 days of sucrose overconsumption. ****$P < 0.0001$. Data are mean ± s.e.m. See Supplementary Table 6 for further details of statistical analyses.

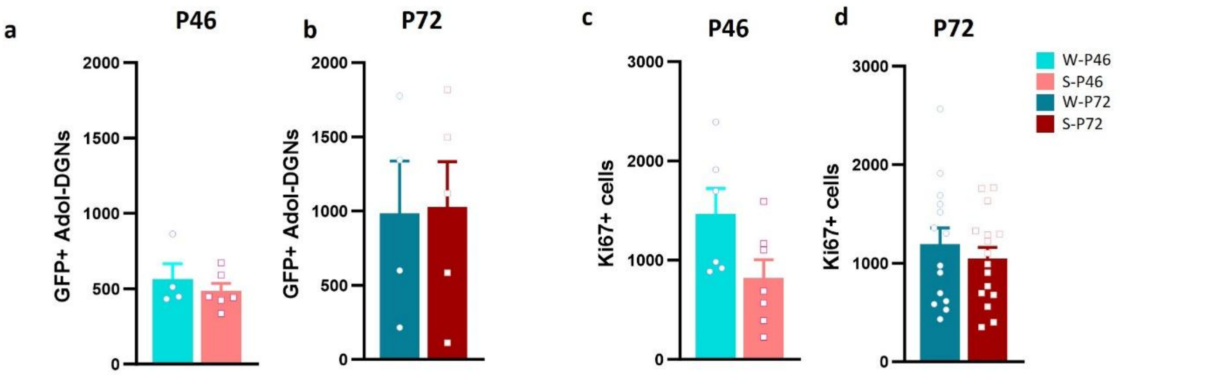


**Supplementary Fig.4: Sucrose overconsumption during adolescence does not alter the number of Adol-DGNs tag with the M-RV GFP (a,b) and Ki67 (c,d).**

Quantification of GFP+ DGNs at P46 (a) and P72 (b) show similar number of cells in water or sucrose animals. The same results were observed when Ki67+ cells were counted at P46 (c) and P72 (d). See Supplementary Table 8 for further details of statistical analyses.

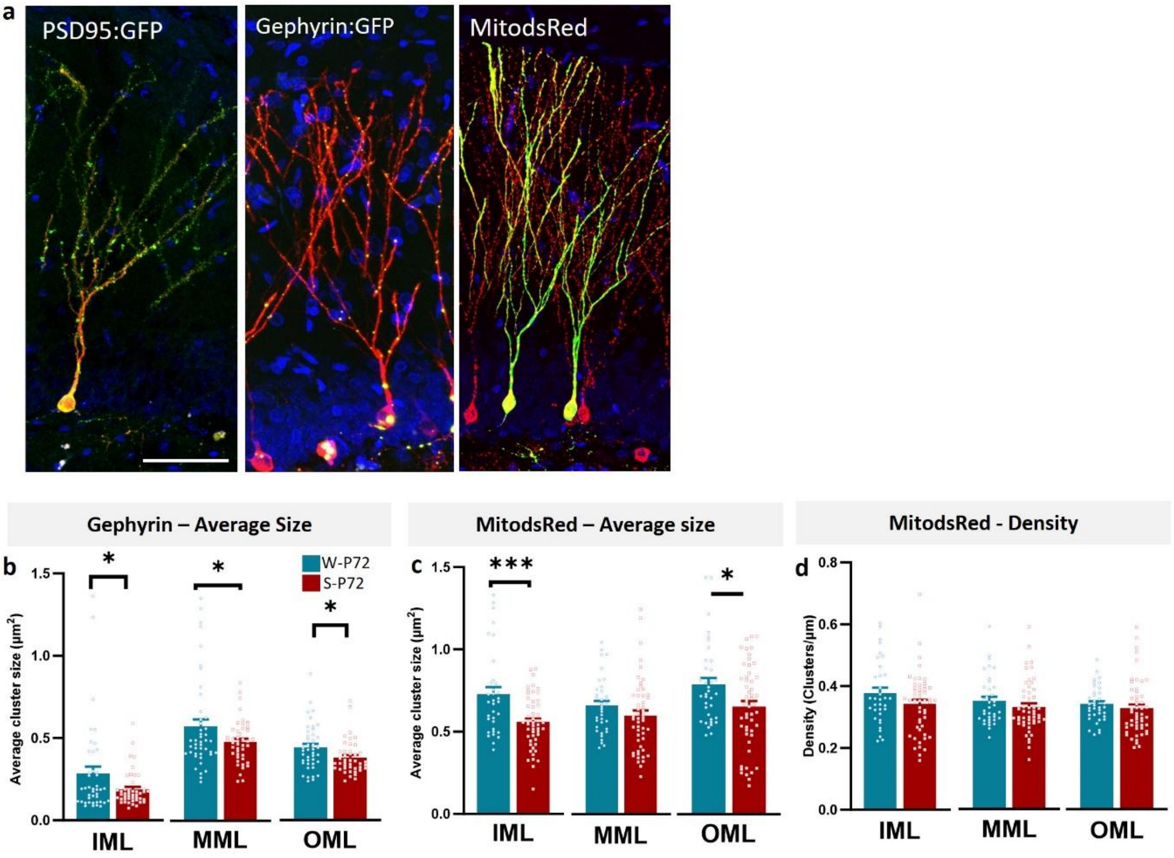


**Supplementary Fig. 5: Sucrose overconsumption during adolescence alters gephyrin and mitochondria average size in the dendrites of Adol-DGNs.**

a. Representative of high-magnification images of Adol-DGNs expressing PSD95:GFP, Gephyrin:GFP and MitodsRed signal at P72. Scale Bar= 50µm. Average size of gephyrin clusters (b) shows a reduction throughout all ML sublayers due to sucrose overconsumption. Same reduction was observed when the average size of mitochondrial clusters was analyzed (c), only in the IML and OML. No significant effects were present in the density of mitochondrial clusters (d). (Gephyrin: n=42-45,44-44,42-43 segments from IML, MML and OML from 4 and 4 animals for W-P72 and S-P72 groups respectively; mitodsRed: n= 34-50, 36-51, 36-52 segments from IML, MML and OML from 4 and 5 animals for W-P72 and S-P72 groups respectively). *$P < 0.05$, ***$P < 0.001$. Data are mean ± s.e.m. See Supplementary Table 10 for further details of statistical analyses.

**Supplementary Table 1: Statistical analyses of Adol-DGNs inhibition during operant behavior for saccharine**

| Group | Nb of rats | FR8 Kinetics | FR8 Lever presses 15min |
|---|---|---|---|
| Water | 10 | TimexCNO: $F_{5,45}$=1.69, p=0.15<br>Time: $F_{5,45}$=16.36, p<0.0001<br>CNO: $F_{1,9}$=3.10, p=0.11 | Paired: $t_9$=2.57, p=0.03<br>One sample: $t_9$=2.57, p=0.03 |

**Supplementary Table 2: Statistical analyses of sucrose overconsumption on different schedule of operant behavior for saccharine**

| Group | Nb of rats | FR1 | FR3 | FR5 | FR8 |
|---|---|---|---|---|---|
| Water<br>Sucrose | 9<br>9 | DayxSuc: $F_{4,64}$=0.80, p=0.52<br>Day: $F_{4,64}$=18.82, p<0.0001<br>Sucrose: $F_{1,16}$=2.39, p=0.14 | DayxSuc: $F_{3,48}$=0.67, p=0.57<br>Day: $F_{3,48}$=18.82, p<0.0001<br>Sucrose: $F_{1,16}$=0.86, p=0.36 | DayxSuc: $F_{5,80}$=0.85, p=0.51<br>Day: $F_{5,80}$=2.66, p=0.02<br>Sucrose: $F_{1,16}$=1.38, p=0.25 | DayxSuc: $F_{9,144}$=0.55, p=0.83<br>Day: $F_{9,144}$=4.16, p<0.0001<br>Sucrose: $F_{1,16}$=4.657, p=0.04 |

**Supplementary Table 3: Statistical analyses of Adol-DGNs activation during operant behavior for saccharine**

| Schedule | Group | Nb of rats | Kinetics | Total Number of Lever presses | Increase LP |
|---|---|---|---|---|---|
| FR8 | Water<br>Sucrose | 9<br>9 | Time: $F_{2,32}$=22.61, p<0.0001<br>CNO: $F_{1,16}$=10.67, p=0.004<br>Sucrose: $F_{1,16}$=4.24, p=0.05<br>TimexCNO: $F_{2,32}$=2.05, p=0.14<br>TimexSuc: $F_{2,32}$=0.84, p=0.44<br>CNOxSuc: $F_{1,16}$=7.18, p=0.01<br>TimexCNOxSuc: $F_{2,32}$=1.64, p=0.20 | CNOxSuc: $F_{1,16}$=7.18, p=0.01<br>CNO: $F_{1,16}$=10.67, p=0.004<br>Sucrose: $F_{1,16}$=4.24, p=0.05 | Unpaired: $t_{16}$=2.68, p=0.01<br>One sample W: $t_8$=3.22, p=0.01<br>One sample S: $t_8$=0.75, p=0.47 |
| PR3 | Water<br>Sucrose | 9<br>9 | Time: $F_{2,32}$=71.54, p<0.0001<br>CNO: $F_{1,16}$=6.42, p=0.02<br>Sucrose: $F_{1,16}$=4.35, p=0.05<br>TimexCNO: $F_{2,32}$=6.47, p=0.004<br>TimexSuc: $F_{2,32}$=6.37, p=0.004<br>CNOxSuc: $F_{1,16}$=0.0001, p=0.99<br>TimexCNOxSuc: $F_{2,32}$=3.17, p=0.05 | CNOxSuc: $F_{1,16}$=2.39, p=0.14<br>CNO: $F_{1,16}$=12.45, p=0.002<br>Sucrose: $F_{1,16}$=6.98, p=0.01 | Unpaired: $t_{16}$=1.54, p=0.14<br>One sample W: $t_8$=3.46, p=0.008<br>One sample S: $t_8$=1.45, p=0.18 |

**Supplementary Table 4: Statistical analysis of Adol-DGNs Erg1 expression after CNO administration**

| Group | Nb of rats | Nb Adol-DGNs | %GFP-Erg1 Colocalization | Erg1+/GCL surface |
|---|---|---|---|---|
| Water | 9 | 118-134 | $t_{16}=2.12, p=0.04$ | $t_{16}=0.93, p=0.36$ |
| Sucrose | 9 | 119-131 | | |

**Supplementary Table 5: Statistical analysis of sucrose overconsumption on dendritic morphology of Gs-DREADD expressing Adol-DGNs**

| Group | Nb of rats | Nb Adol-DGNs | T Dendritic Length | Nb of Ends | Sholl Analysis |
|---|---|---|---|---|---|
| Water | 9 | 54 | $t_{103}=2.46, p=0.01$ | $t_{103}=1.25, p=0.21$ | DistxSuc: $F_{35,3605}=4.11, p<0.0001$ |
| Sucrose | 9 | 51 | | | Distance: $F_{35,3605}=137.6, p<0.0001$ |
| | | | | | Sucrose: $F_{1,103}=9.47, p=0.002$ |

**Supplementary Table 6: Fluid intake during sucrose overconsumption**

| Sucrose effect | Day effect | Sucrose x Day effect |
|---|---|---|
| $F_{1,16}=51.50, p<0.0001$ | $F_{6,96}=2.46, p=0.02$ | $F_{6,96}=1.01, p=0.41$ |

**Supplementary Table 7: Total number of Adol-DGNs infected in all the experiments (methods section)**

| Group | Gi-DGNs | Gs-DGNs | P46-GFP | P72-GFP | P72-PSD95 | P72-Gephyrin |
|---|---|---|---|---|---|---|
| Water | 3538±308 | 4214±715 | 564±101 | 984±352 | 452±82 | 808±387 |
| Sucrose | - | 3918±489 | 485±50 | 1026±307 | 537±83 | 434±215 |

**Supplementary Table 8: Statistical analysis of GFP+ DGNs and Ki67+ cells at P46 and P72 after sucrose overconsumption**

| Group | P46 GFP+ DGNs | P72 GFP+ DGNs | P46 Ki67+ cells | P72 Ki67+ cells |
|---|---|---|---|---|
| Water<br>Sucrose | $t_8$=0.77, p=0.46 | $t_7$=0.93, p=0.08 | $t_{11}$=2.09, p=0.06 | $t_{28}$=0.73, p=0.46 |

**Supplementary Table 9: Statistical analysis of sucrose overconsumption in dendritic morphology of Adol-DGNs**

| Delay | Group | Nb of rats | Nb Adol- DGNs | T Dendritic Length | Nb of Ends | Sholl Analysis |
|---|---|---|---|---|---|---|
| P46 | Water | 4 | 33 | $t_{79}$=1.50, p=0.13 | $t_{79}$=1.35, p=0.17 | DistxSuc: $F_{31,2449}$=0.74, p=0.84 |
| | Sucrose | 6 | 48 | | | Distance: $F_{31,2449}$=94.14, p<0.0001 |
| | | | | | | Sucrose: $F_{1,79}$=2.45, p=0.12 |
| P72 | Water | 4 | 32 | $t_{72}$=3.407, p=0.001 | $t_{72}$=2.172, p=0.03 | DistxSuc: $F_{31,2232}$=5.33, p<0.0001 |
| | Sucrose | 5 | 42 | | | Distance: $F_{31,2232}$=130.8, p<0.0001 |
| | | | | | | Sucrose: $F_{1,72}$=9.09, p=0.003 |

**Supplementary Table 10: Statistical analysis of sucrose overconsumption on connectivity and mitochondrial network of Adol-DGNs**

| Virus | Group | Nb of rats | Nb segments | Parameter | IML | MML | OML |
|---|---|---|---|---|---|---|---|
| PSD95:GFP | Water | 5 | 68/69/65 | Density | $t_{169}$=0.01, p=0.98 | $t_{172}$=0.59, p=0.55 | $t_{167}$=2.84, p=0.005 |
| | Sucrose | 8 | 103/105/104 | | | | |
| Gephyrin:GFP | Water | 4 | 42/44/42 | Density | $t_{85}$=2.99, p=0.003 | $t_{86}$=0.64, p=0.52 | $t_{83}$=0.11, p=0.90 |
| | Sucrose | 4 | 45/44/43 | Av Size | $t_{85}$=2.15, p=0.03 | $t_{86}$=2.02, p=0.04 | $t_{83}$=2.36, p=0.02 |
| MitodsRed | Water | 4 | 34/36/36 | Total Absolute Area | $t_{82}$=3.76, p=0.0003 | $t_{85}$=2.57, p=0.01 | $t_{86}$=3.63, p=0.0005 |
| | Sucrose | 5 | 50/51/52 | Density | $t_{82}$=1.58, p=0.11 | $t_{85}$=1.10, p=0.27 | $t_{86}$=0.76, p=0.44 |
| | | | | Av Size | $t_{82}$=3.69, p=0.0004 | $t_{85}$=1.38, p=0.16 | $t_{86}$=2.48, p=0.01 |
